# Forward planning in a population-based alcohol use disorder sample

**DOI:** 10.1101/2022.11.21.517329

**Authors:** Johannes Steffen, Pascale C. Fischbach, Lorenz Gönner, Stefan J. Kiebel, Michael N. Smolka

**Author notes:** Corresponding author Corresponding author Michael N. Smolka Full postal address: Section of Systems Neuroscience Department of Psychiatry Technische Universität Dresden Würzburger Str. 35 01187 Dresden (Germany).

## Abstract

**Background:** Etiological theories of addictive behavior postulate a key role for decision-making mechanisms. However, current research is lacking clear computational models for decision-making in multi-step forward planning scenarios to identify underlying mechanisms and make derived hypotheses testable.

**Methods:** We used a recently developed planning task and computational model to investigate performance, planning time and inferred forward planning parameters like planning depth and decision noise in 30 individuals diagnosed with mostly mild-to-moderate alcohol use disorder (AUD) relative to 32 healthy control participants, both sampled from the general population.

**Results:** Contrary to our hypothesis, we did not observe reduced planning depth in participants with AUD but found that participants with AUD showed a higher performance in the planning task. Group differences could be explained by planning time and general cognitive performance. Importantly, participants with AUD invested more time for planning, showed a higher correlation of planning depth with incentive value and showed lower response noise, potentially indicative of higher choice consistency.

**Conclusion:** The significant differences in planning time, moderation of planning depth by incentive value and choice consistency may reflect higher motivation and willingness to exert effort among participants with AUD compared to healthy controls. Overall, our findings do not support the notion that mild-to-moderate alcohol use disorder is associated with impairments in forward planning across multiple steps.

## INTRODUCTION

Substance use disorders frequently impair physical and mental health and have serious social and economic consequences. Especially the harmful use of alcohol is one of the most important risk factors for population health worldwide (World Health Organization, 2018). Despite these detrimental effects, our understanding of the mechanisms and trajectories of alcohol use disorders (AUD) remains limited. According to the “transition-to-habit” theory (Everitt & Robbins, 2016), the process of developing an AUD begins with reflective goal-directed decisions for consuming alcohol. Here, individuals might underestimate the distal risks of alcohol use because they do not adequately evaluate future consequences (MacKillop et al., 2011; Story et al., 2014) leading to repetitive consumption and a development of ‘bad’ alcohol use habits. The exact computational principles of this adverse decision-making of certain individuals at this early stage are still an unresolved issue. However, identifying such a computational principle is important to derive predictors for long-term problematic alcohol use. One plausible computational approach is model-based forward planning, which rests upon knowledge, represented as a probabilistic model, about the potential consequences in the future and a weighing of their negative and positive values for making decisions (Dolan & Dayan, 2013; Sutton & Barto, 2018). One obvious challenge of model-based forward planning is the question of how deep, i.e. how far into the future, one should plan ahead (Steffen et al., 2022). Deeper forward planning potentially leads to more accurate valuations of consequences but is also computationally more costly resulting in a complex cost-benefit tradeoff where resource-rational solutions are not obvious. Regarding decisions on alcohol consumption however, positive effects typically occur quickly after consumption while negative consequences manifest at a later stage and usually over sequences of further actions. Hence, forward planning should be a crucial mechanism in this context where higher planning depths could be critical for foreseeing the (re-)occurrence of problematic outcomes and to therefore disrupt the development of habitual alcohol use. In this study, we aimed at experimentally testing the hypothesis that individuals with AUD show a lower planning depth. By comparing healthy controls (HC) to general-population non-treatment-seeking individuals fulfilling an AUD diagnosis, we specifically targeted a stage of AUD with still acceptable levels of functioning.

So far, we are not aware of previous studies examining planning depth in AUD. However, related evidence comes from two branches of research: First, planning capabilities have classically been studied using sequential problem-solving tasks like the Tower of London Task (ToL; Shallice, 1982). While these tasks do not directly assess planning depth, they provide indicators of forward planning performance, where lower performance usually means more moves needed to reach a target configuration. In a meta-analysis of 39 studies by Stephan et al. (2017), clinical AUD samples showed a significantly lower planning performance according to a planning and problem-solving composite score. Therefore, forward planning capabilities are clearly associated with AUD. However, clinical studies assessing recently detoxified patients should be reviewed with caution as planning could similarly be impaired due to toxic effects of long-time alcohol consumption on frontal lobe brain tissue (Chanraud et al., 2007).

The second branch of research has been using the two-stage Markov task by Daw et al. (2011) to study the balance between model-based and model-free control in sequential decision-making, where the former is equivalent to a form of forward planning. Studies using this approach in clinical AUD samples either found no difference in model-based control compared to healthy controls (Sebold et al., 2017; Voon et al., 2015) or the effect did not survive control for performance in a neuropsychological test (Sebold et al., 2014). However, Voon et al. (2015) found a positive association of the model-based weight parameter with the number of weeks participants were abstinent indicating that forward planning does play a role in subsequent control of consumption behavior. Sebold et al. (2017) found a similar association of model-based control and 2-year relapse rates specifically in patients with high self-reported drinking expectations at baseline. Studies focusing on general-population samples showed a similar picture. While two studies focusing on lifetime drinking patterns of 18-year-olds (Nebe et al., 2018) or undergraduate students (Byrne et al., 2019) did not find robust associations with model-based control, we suspect that their specific operationalizations captured experimentation with alcohol during adolescence rather than pathological use (e.g. age of first drink). In contrast, Gillan et al. (2016) did find a significant negative association of the number of self-reported AUD symptoms with model-based control in a large general-population sample. Moreover, a longitudinal assessment of the sample of Nebe et al. (2018) revealed that model-based control predicted binge drinking trajectories for the subsequent years (Chen et al., 2021). Similarly, Ebrahimi et al. (2024) showed an association of higher model-based control with more successful implementation of intentions to reduce drinking during a 12-month follow-up ecological ambulatory assessment.

Taken together, previous studies showed that forward planning is associated with alcohol-related outcomes. Planning seems to be particularly predictive for future drinking behavior e.g. by preventing risky binge drinking trajectories (Chen et al., 2021) or promoting intended reduction (Ebrahimi et al., 2024) or abstinence (Sebold et al., 2017; Voon et al., 2015). However, these approaches did not yet allow to differentiate computational parameters of forward planning like planning depth or a general response bias or response noise, which may enable more detailed insights into the cognitive mechanisms underlying interindividual differences.

To assess planning depth in the described sample in comparison to HC, we used a sequential decision-making task, the Space Adventure Task (SAT), as described already by Steffen et al. (2022). Participants’ choices in this task were modelled with a reinforcement learning agent model with planning depth as a free parameter that could be inferred with Bayesian inference (see Methods section).

Forward planning capabilities are assumed to be correlated with several cognitive parameters like working memory, processing speed and general crystalline intelligence (Otto et al., 2013, 2014; Sebold et al., 2014; Steffen et al., 2022). These have been shown to be impaired in AUD samples (Chanraud et al., 2007; Sebold et al., 2014; Stephan et al., 2017). We therefore considered measures of these constructs as covariates in our analysis to investigate to what extent they might explain differences in planning depth.

Based on previous research, we hypothesized that in the SAT, participants fulfilling an AUD diagnosis compared to HC should demonstrate reduced forward planning capabilities indicated by lower scores and a lower inferred planning depth. We also expected performance measures for the three assessed general cognitive parameters to be positively associated with planning depth while not fully explaining group differences in planning depth.

## MATERIALS AND METHODS

### Participants and procedure

Potential candidates were recruited from the Dresden city area by distribution of flyers and online advertisement which specifically targeted people who drink alcohol regularly and/or in large amounts. Interested candidates were pre-screened via telephone for our inclusion and exclusion criteria. We included individuals aged from 16 to 65 years. Fulfilment of the criteria of AUD during the past 12 months according to the fifth edition of the Diagnostic and Statistical Manual of Mental Disorders (DSM-5; American Psychiatric Association, 2013) were diagnosed with a structured clinical interview (SCID-5-CV; First, 2016). Individuals were excluded if they reported any physical withdrawal symptoms from alcohol, a lifetime history of mania or psychosis, an acute severe major depression episode or suicidal thoughts, a severe illness of the central nervous system or any previous or current substance dependencies other than alcohol or tobacco. Female participants were additionally excluded in case of pregnancy or if they were currently nursing an infant. On site, participants were only assessed if they had a measured breath alcohol value of 0% and no positive urine test result for any substance other than cannabis. After giving written consent, participants filled out a sociodemographic questionnaire, worked on the SAT as well as the neuropsychological tests and finally, completed questionnaires on quantity and frequency of alcohol-, tobacco- and cannabis use. At the end, participants reporting 2 or more AUD symptoms were seen by our study physician and offered contact with the outpatient clinic or addiction counseling centers. The overall procedure took approximately 2.5-3 hours and participants were compensated with 25-35€ depending on their performance in the SAT. The ethics committee of the TU Dresden gave ethical approval for this work (EK514122018).

Out of 70 invited individuals, 5 dropped out of the assessment for several reasons: they did not show up (*n =* 1), they only showed up intoxicated (*n =* 1), technical issues during the task (*n =* 2) and exclusion because the control group sample was already complete (*n =* 1). Out of the resulting 65 completed assessments, three more participants had to be excluded due to behavior in the main task of the experiment that was close to random. The remaining *N =* 62 participants were either assigned to the AUD group (*n =* 30) fulfilling 2 or more AUD criteria or to the HC group (*n =* 32) fulfilling less than 2 AUD criteria. According to the DSM-5 degrees of AUD severity, the great majority (90%) of the AUD group showed a mild to moderate form of AUD (see Figure S1). In the alcohol use disorder identification test (AUDIT; Saunders et al., 1993), the AUD group scored significantly higher (*t =* -6.702, *p <* .001) with mean values and standard deviation 14.07 ± 4.4 for the AUD group and 6.44 ± 4.6 for the HC group. The two groups did not show significant differences in education or employment status, while participants of the AUD group did report higher frequencies of heavy episodic drinking and higher smoking quantities (see Table 1). For a detailed depiction of quantity and frequency of participants’ tobacco- and alcohol use see Figures S2-S7.

**Table 1.** Descriptive Statistics and group comparison of task outcomes and model parameters.

|  | AUD<br>(N = 30) | HC<br>(N = 32) | *t* | *p* |
| --- | --- | --- | --- | --- |
| **Sample Characteristics** |  |  |  |  |
| Age | 28.3 (8.6) | 29.8 (10.8) | -.05<sup>a</sup> | .961 |
| Gender (Male/Female) | 18/12 | 13/19 | 2.33<sup>b</sup> | .127 |
| DSM-5 AUD Criteria<br>(no/mild/moderate/severe) | 0/18/9/3 | 32/0/0/0 | - | - |
| AUDIT | 14.07 (4.4) | 6.44 (4.6) | -6.70*** | <.001 |
| HED frequency last 30 days <sup>c</sup> | "6-9 days" <sup>c</sup> | "0 days" <sup>c</sup> | 24.71*** <sup>b</sup> | <.001 |
| Smoking quantity last 30 days <sup>d</sup> | "< 1/day" <sup>d</sup> | "0" <sup>d</sup> | 16.58* <sup>b</sup> | .011 |
| HEEQ (Yes/No) | 25/5 | 26/5<sup>e</sup> | .34<sup>b</sup> | .561 |
| Employed (Yes/No) | 25/5 | 25/6<sup>e</sup> | 1.32<sup>b</sup> | .251 |
| **Model Parameters** |  |  |  |  |
| Mean Planning Depth | 2.23 (.16) | 2.11 (.26) | 1.96<sup>a</sup> | .05 |
| Learning Rate $\alpha$ | .00 (.00) | .04 (.04) | -6.15*** | <.001 |
| Inverse Response Noise $\beta$ | 2.14 (.37) | 1.72 (.74) | 2.82*** | <.001 |
| Response Bias $\theta$ | -.35 (.43) | -.11 (.42) | -2.17* | .034 |
| **Task Performances [%]** |  |  |  |  |
| Space Adventure | 68.6 (9.9) | 58.2 (15.4) | 3.18** | .002 |
| Identical Pictures | 65.1 (9.3) | 62.6 (10.5) | 1.02 | .310 |
| Spatial Working Memory | 94.1 (4.9) | 90.4 (7.1) | 2.41* | .019 |
| Raven's Matrices | 67.8 (18.9) | 59.4 (21.8) | 1.89<sup>a</sup> | .058 |
| **Task Reaction times [s]** |  |  |  |  |
| Space Adventure | 9.9 (5.0) | 7.2 (3.9) | 2.41* | .019 |
| Identical Pictures | 2.4 (.5) | 2.5 (.5) | -1.17<sup>a</sup> | .242 |
| Raven's Matrices | - | - | - | - |
| Spatial Working Memory | 1.2 (.2) | 1.3 (.3) | -1.98<sup>a*</sup> | .047 |

**Note.** Descriptive statistics are formatted as: Mean (Standard Deviation).
HEEQ = German Higher Education Entrance Qualification.
AUDIT = Alcohol Use Disorders Identification Test.
HED = heavy episodic drinking.
<sup>a</sup> Z-statistic and corresponding asymptotic p-value: because of non-normality, an additional non-parametric Mann-Whitney-U test was performed which yielded different results.
<sup>b</sup> Pearson's $\chi^2$ -statistic and corresponding p-value.
<sup>c</sup> Median questionnaire response for heavy episodic alcohol drinking frequency of last 30 days (see Figure S4).
<sup>d</sup> Median questionnaire response for tobacco smoking quantity of last 30 days (see Figure S6).
<sup>e</sup> Data missing for one HC participant due to technical failure.

### Neuropsychological Tests

We used the first subtask of the computerized Spatial Working Memory (SWM) task (Nagel et al., 2008) as a measure for spatial processing abilities. In this task, participants are presented with a 4 x 4 grid of circles in which a series of dots is displayed consecutively in specific locations of the grid. At the end of each sequence one circle is marked and participants have to indicate whether a dot was presented at that position or not. As a measure of processing speed, participants completed the Identical Pictures Task (IDP; Lindenberger et al., 1993). In this task, participants are presented with five symbols and must quickly and accurately find the symbol that matches the one at the top. Finally, as a measure of fluid intelligence, we used a computerized implementation of the short-form of Raven’s Advanced Progressive Matrices (RAV; Bors & Stokes, 1998). For the SWM, the IDP and the RAV, we acquired reaction time (RT) and accuracy and excluded trials with an RT below 200 ms. As a measure of performance, we computed the achieved percentage of maximum possible correct responses. For a more detailed description of the neuropsychological tests, see Figure S8-S10.

### Space Adventure Task

The SAT is a computerized sequential decision-making task which requires forward planning to be performed effectively. During a fixed sequence of 100 mini-blocks, participants visited planets with their spaceship and had to accumulate as much fuel as possible (indicated by a fuel bar at the top). Figure 1A shows the planet types with their respective fuel rewards. Figure 1B depicts an example mini-block with three steps left (green squares) and low noise (black background). The mini-blocks were designed in a way that participants had to plan forward to find the route leading to maximum fuel gain with either two or three steps. At each step, participants could choose between two actions: (i) move to the next planet in a clockwise fashion by using 2 fuel points or (ii) jump to a non-neighboring planet as determined by a given travel pattern (Figure 1C) which consumed 5 fuel points. While moving clockwise was deterministic and always brought one to the target planet, jumping was assigned a certain probability for the transition to fail causing the spaceship to land instead on one of the two neighboring planets of the target planet, each with equal probability. The probability that jumping would succeed was varied block-wise between 50% and 90% (‘noise condition’). However, as pointed out in Steffen et al. (2022), this condition was confounded with the training effect and with the maximum possible reward. Therefore, we refrain from analyzing this condition here. Prior to the main set of mini-blocks, participants were explicitly instructed about all aspects of the tasks, implicitly learned the jumping failure probability by trial and error, memorized the travel pattern and familiarized themselves with the procedure during 20 training mini-blocks. The SAT was implemented in MathWorks MATLAB version 2018a and run on a standard PC. Participants entered their responses via keyboard with key ‘S’ (in German ‘Sprung’) indicating the jump action and the right arrow key indicating the move action. For a detailed description of the SAT see Steffen et al. (2022).

**Figure 1.**
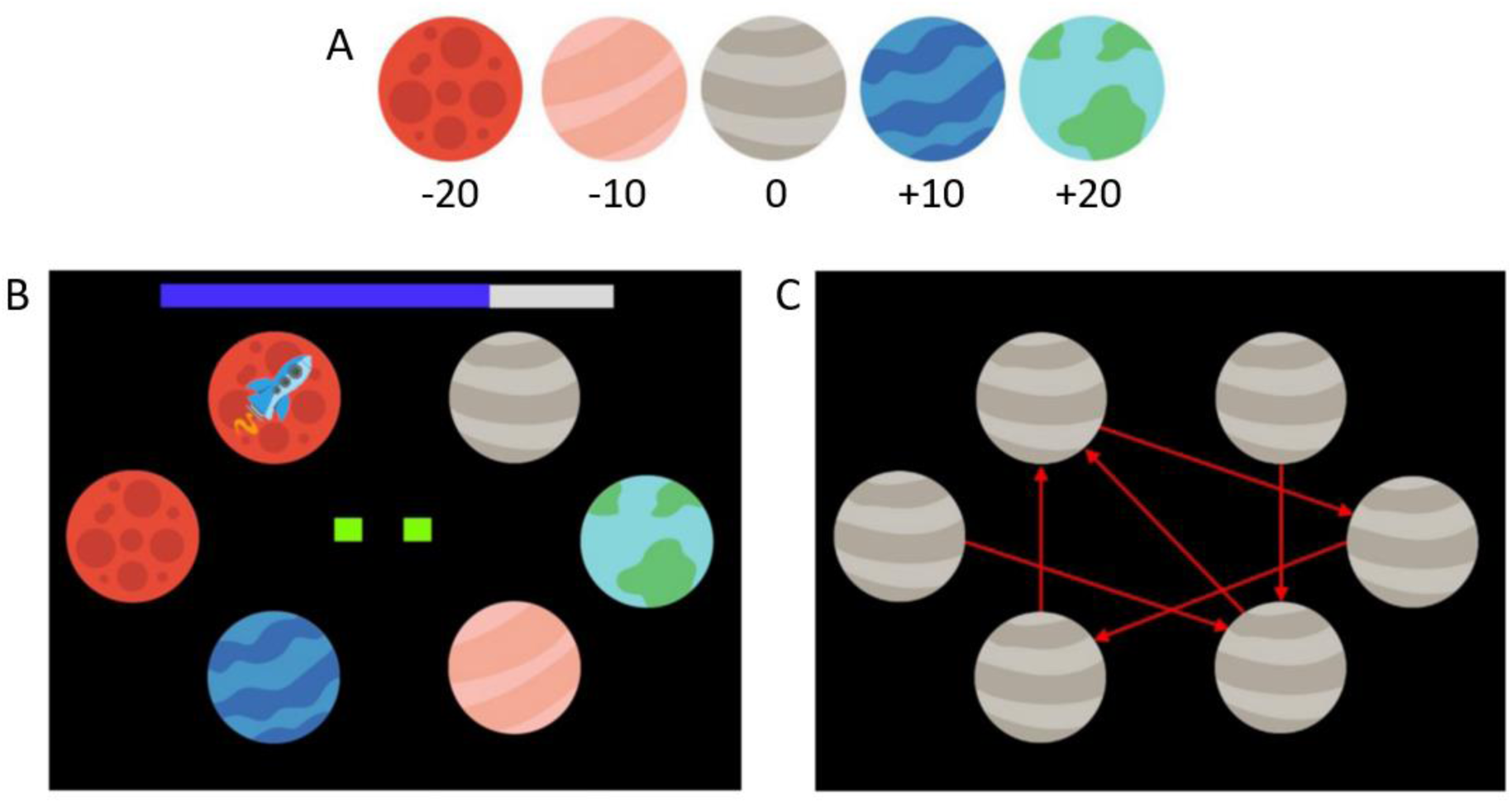
Schematic of the Space Adventure Task. **(A)** Five planet types with different amounts of fuel to win or lose when landing on them. **(B)** Planet configuration in a specific mini-block with rocket on starting planet. On top of the screen, current fuel is indicated by the blue part of the bar. The two green squares in the middle of the screen show that two steps are left. The black background without asteroids indicates high jumping success probability. **(C)** The jump pattern which was not visible during the experiment and had to be memorized before.

### Computational model

We modelled participants’ action choices with a mixture model of three single model-based RL agent models with planning depth of 1, 2 and 3, respectively. Each agent had an optimal model of the environment, i.e. it was completely informed about the rules of the task. This environment model entailed the set of available actions 𝐴 = {′𝑚𝑜𝑣𝑒^′^, ^′^𝑗𝑢𝑚𝑝^′^} and states 𝑆 (the planet configuration of the mini-block), the transition probabilities 𝑝(𝑠_𝑡+1_|𝑠_𝑡_, 𝑎_𝑡_) for reaching a subsequent state 𝑠_𝑡+1_ from a given state 𝑠_𝑡_ with action 𝑎_𝑡_ as well as the immediate reward of reaching each state 𝑟(s) indicated by the planet types of the current configuration. To choose the optimal action for a specific state in a specific mini-block, the agents computed the expected cumulative reward for executing each action with an optimal forward planning algorithm (value iteration) only limited by their planning depth.

Value iteration outputs the expected cumulative reward for executing each action 𝑎 in the current state 𝑠 with planning horizon 𝑑, which is called the state-action- or 𝑄-values. These 𝑄-values were the essential values for action selection. The higher the relative value of an action was, the higher should have been the probability of selecting that action. Action selection was therefore modelled probabilistically with a softmax function (Sutton & Barto, 2018). For our case of two available actions, this corresponded to a sigmoid transformation 𝜎(𝑥) of the difference between the 𝑄-values, Δ𝑄(𝑠_𝑡_, 𝑑). Choice probabilities were thus defined as:

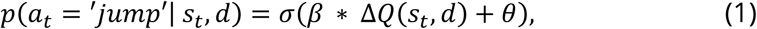

where

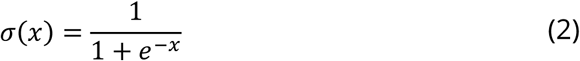

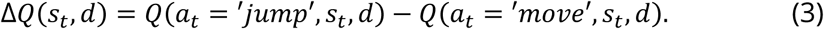

Here, the inverse response noise *β* controlled the extent to which differences in 𝑄-values affected action selection. If *β* = 0, actions are selected with a constant probability independent of outcomes, while higher values of *β* represented higher probability to select the action with the highest 𝑄-value. The parameter 𝜃 denoted an a priori response bias, where positive values imply a bias towards choosing ’jump’.

Since the state transition probabilities for the jump action 𝑝(𝑠_𝑡+1_| 𝑠_𝑡_, 𝑎_𝑡_ = ′𝑗𝑢𝑚𝑝′) were not given explicitly during training, we assumed an experience-based learning process of the state transition probabilities. The belief about the probability that a jump will be successful 𝜌 = 𝑝(𝑠_𝑡+1_ = ′𝑡𝑎𝑟𝑔𝑒𝑡′|𝑠_𝑡_, 𝑎_𝑡_ = ′𝑗𝑢𝑚𝑝′) was updated using the temporal difference rule:

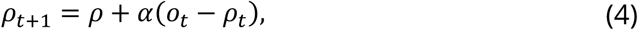

depending on the experienced success (𝑜_𝑡_ = 1) or failure (𝑜_𝑡_ = 0) of a jump. The learning rate parameter 𝛼 modelled how fast participants changed their beliefs about the probability of transition success. Larger values of 𝛼 could also be interpreted as faster forgetting of prior experience and stronger reliance on recent outcomes.

### Planning depth and parameter inference

To infer the four free model parameters, inverse response noise *β*, response bias 𝜃, learning rate 𝛼 and planning depth 𝑑, from participants’ choices, we used a hierarchical probabilistic model of free parameters and approximate Bayesian inference scheme. The approximate inference can be imagined as a two-step procedure. Firstly, approximate posteriors of *β*, 𝜃 and 𝛼 were computed for each of the three RL agent models with planning depth 𝑑 ∈ {1,2,3}. Secondly, these agents with their respective inferred parameter distributions for inverse response noise *β*, response bias 𝜃 and learning rate 𝛼 were used to infer the posterior over planning depth 𝑑. For this purpose, the response likelihood of each participant was modelled as a mixture model of response likelihoods of these three agents, which can be expressed as:

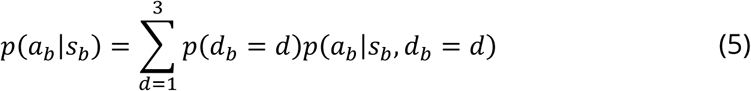

Importantly, we assumed that forward planning should mostly happen before the first action of each mini-block. Hence, for the model inversion (parameter inference), we have constrained behavioral data only to the first choice in each mini-block.

In Equation (5) defining the mixture model, 𝑝(𝑑_𝑏_ = 𝑑) denotes the prior probability each planning depth has in mini-block 𝑏. To account for the limited amount of behavioral data, and for expected within-subject similarities in participants’ responses, we designed the probabilistic generative model in a hierarchical fashion with a participant-level and mini-block-level dependence of parameter values. The parameters *β*, 𝜃 and 𝛼 were modelled on the participant-level, while planning depth 𝑑 was modelled on the mini-block-level. As an analytical solution for the posteriors of the parameters was intractable, we instead used the stochastic variational inference scheme from the probabilistic programming library Pyro v1.5.2 (Bingham et al., 2019) to approximate the posterior distributions. For more details on the computational model and inference procedure see Steffen et al. (2022).

### Statistical Analysis

Statistical analysis was performed using IBM SPSS Statistics version 28. For all relevant variables and measures (including RT, performance, inferred model parameters, sociodemographic, and cognitive covariates), we tested for normal distribution via the Shapiro-Wilk test, and for variance homogeneity across groups via Levene’s test. If non-normality was a concern, we used the Wilcoxon test for group comparisons. In case of variance inhomogeneity, we compared groups using a permutation test with 1,000 samples. Otherwise, we used two-sided *t*-tests. For categorical data, we performed Pearson χ² tests. For ease of reporting, we only report the results of *t*-tests whenever their result was highly similar to other tests used.

Before analyzing behavioral data, we first checked each dataset for random behavior in the SAT as indicated by the inverse response noise *β*. For this, we ran simulations with multiple *β* values, 1,000 repetitions each, and found that *β* should have at least a value of 1 to be far enough from causing random behavior, i.e. for planning depths to be distinguishable. We excluded three participants with less than 500 total points, equivalent to the mean end result achieved by agents with planning depth 𝑑 = 1 and *β* = 1. We also considered RTs until the first action as criterion for exclusion, which should not be below the common timeframe of 200 ms for perceptual and motor processes (Whelan, 2008). However, no participant had any RT below 200 ms. For the analysis, mean planning depths and RTs were averaged for each participant.

To evaluate mini-block-wise mean planning depth in more detail, we used a linear mixed-effects model with random intercept and random effect for steps condition per participant as well as fixed effects for group and a group-by-condition interaction term (Model 1.1).

Further models were set up to control for covariation of neuropsychological variables with mean planning depth and SAT performance. We included all covariate measure that showed significant 2-sided Pearson correlation with these outcomes. In linear mixed-effects model 1.2 for mean planning depth, we therefore added fixed effects for performance in the SWM and RAV task to model 1.1. To investigate the role of planning time, we also included SAT reaction time until first action as a predictor. We compared the fit of both models using a likelihood-ratio test. To control for covariates in SAT performance, we conducted a linear regression analysis with a group indicator and covariates IDP-, SWM- and RAV task performance as well as SAT reaction time as predictors.

In a post-hoc analysis of motivational differences between groups in the SAT, we first computed Q-values for each mini-block, action and planning depth and extracted the maximum absolute value for each mini-block (Q*) as an indicator for the incentive value of the mini-block. We then analyzed mean planning depths with a linear mixed-effects model with random intercept and random effect for Q* per participant as well as fixed effects for group and a group-by-Q* interaction term (Model 1.3).

## RESULTS

We report descriptive statistics in Table 1. When analyzing for outlier data based on RTs below 200 ms, no SAT mini-blocks had to be excluded. In the SWM task, three participants had one trial with an RT below 200 ms that had to be excluded.

### Performance and planning depth

We found that in the SAT, Participants with AUD showed a higher performance (*t*(53.3) = 3.18, *p =* .002) while taking more time to react (*Z =* 2.41, *p =* .019). There was no significant difference in mean planning depth compared to HC participants (Z = 1.96, p = 0.05; see Table 1 and Figure 2B). Moreover, reaction times were positively associated both with performance (*r =* .57, *p* < .001) and with planning depth (*r =* .49, *p* < .001, see Figure 2C and refer to Figure S13 for a full correlation matrix). A post-hoc linear regression analysis with a “group x reaction time” interaction term revealed that the association with performance was significantly higher in the HC group (*t =* -3.342, *p =* .001; see Table S3). This was not the case for the association with planning depth (95%CI_BCa_ [-.054, .003], *p =* .084; see Table S2). The post-hoc analysis of motivation on planning depth showed a significant positive effect of absolute maximum Q-values (*b =* .018, *t*(76.9) = 16.68, *p <* .001; see Table S4), which varied significantly between participants (*Z =* 2.47, *p =* .014), as well as a significant group-by-Q* interaction (*b =* .004, *t*(76.9) = 2,74, *p =* .008; see Figure S12) potentially indicating higher motivation through the incentive value of the mini-block in the AUD group.

**Figure 2.**
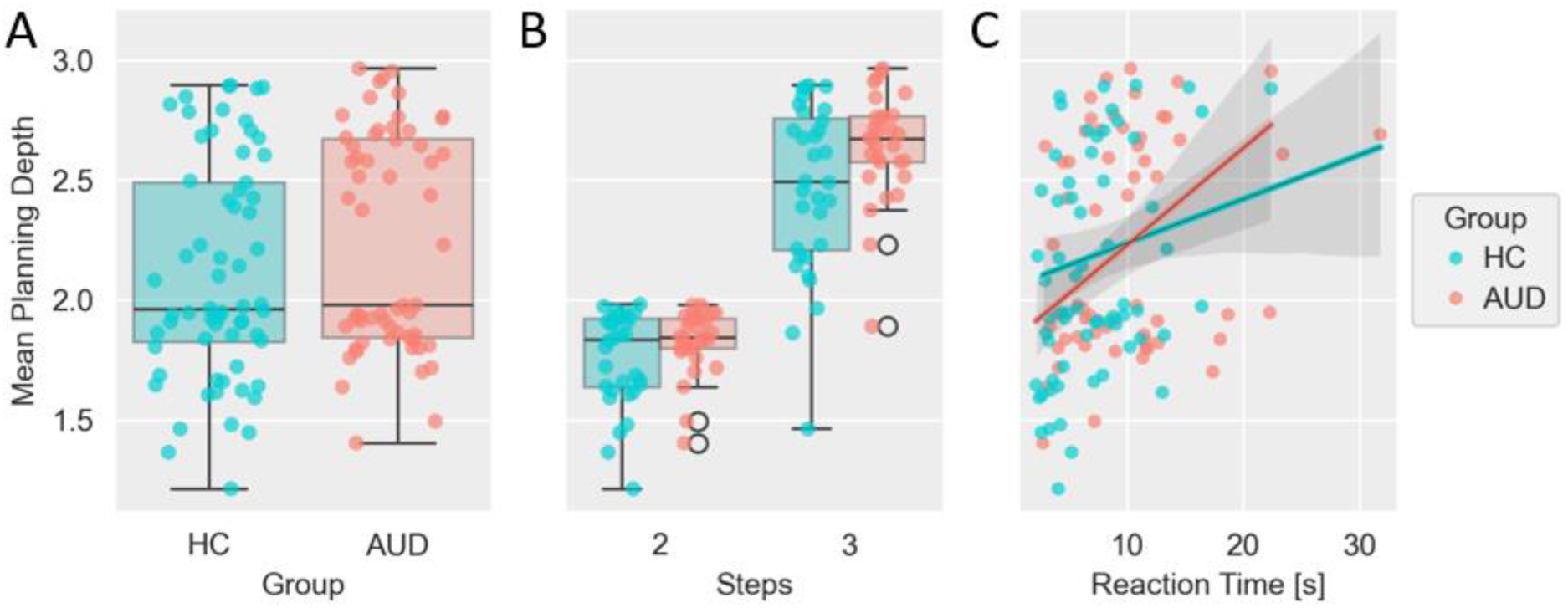
Inferred mean planning depth per participant and correlation with reaction time until first action in the Space Adventure Task. **(A)** Boxplot and scatterplot for mean planning depth averaged over the whole task for each participant comparing healthy controls (HC) and participants with alcohol use disorder (AUD). **(B)** Same plot as (A) with mean planning depth averaged separately for each step condition. **(C)** Scatterplot of mean planning depth and reaction time until first action for each participant and correlation of these two measures for each group.

**Figure 3.**
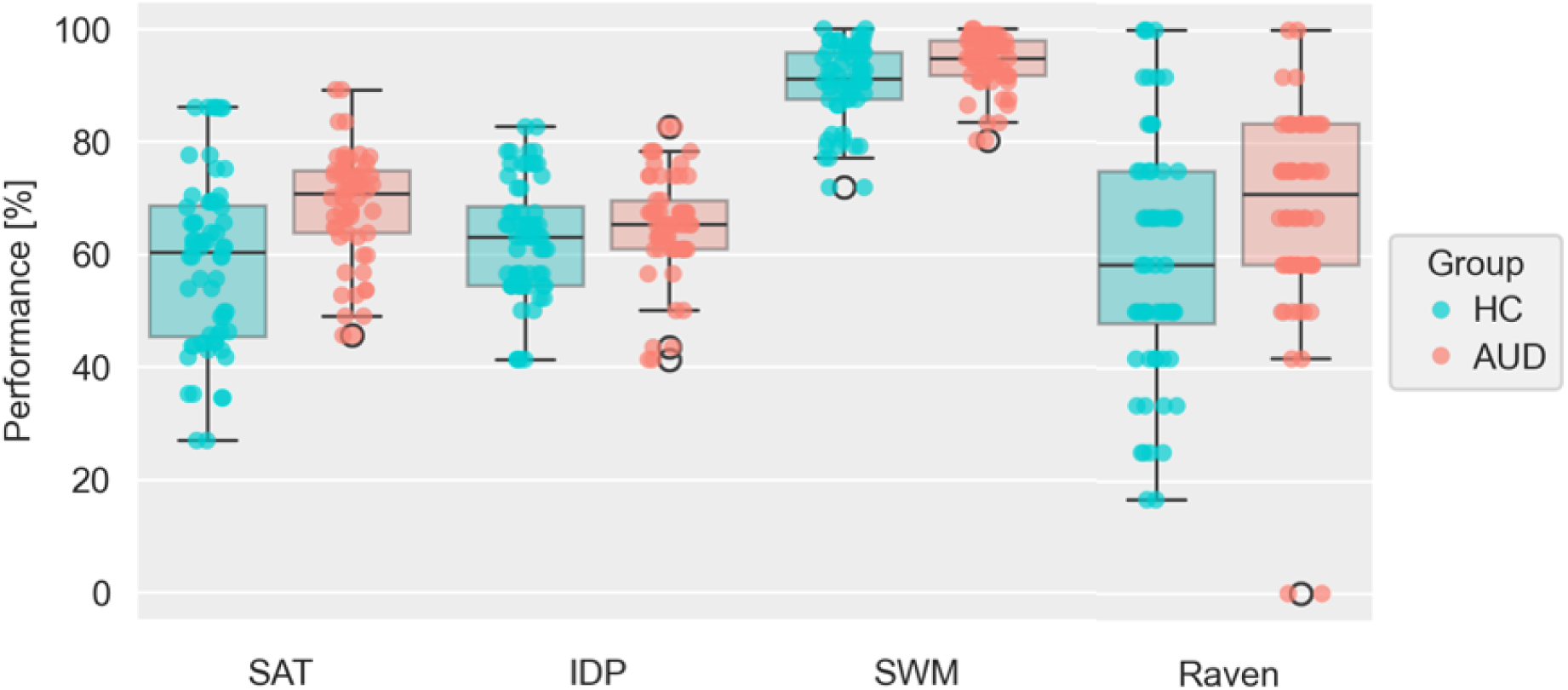
Performances per participant in the Space Adventure Task (SAT) and cognitive covariate tasks. Boxplot and scatterplot of participants’ achieved performance in the SAT, Identical Pictures Task (IDP), Spatial Working Memory Task (SWM) and short-form of Raven’s Advanced Progressive Matrices (Raven) comparing healthy controls (HC) and participants with alcohol use disorder (AUD). Performance was computed as achieved percentage of maximum possible points gained for the SAT and achieved percentage of maximum possible correct responses for the other tasks.

To analyze the effect of the steps condition (i.e. the number of sequential actions to be considered during planning, either two or three) on planning depth, we performed a linear mixed-effects analysis (Model 1.1), which revealed a main effect of steps (*t*(62.3) = 18.39, *p <* .001), but no main effect of group (*t*(62.3) = 1.60, *p =* .114) or group x steps interaction (*t*(62.3) = 1.91, *p =* .061; see Table 2). Within the 2-step condition, most participants’ choices were close to optimal, while mean planning depth increased by approximately 0.7 steps from the 2-step to the 3-step condition (Figure 2B). As indicated by the random effects of the model, there was significant between-subject variation of the intercept (*η*(62.3) = .027, *p <* .001) and of the steps effect (*η*(62.3) = .044, *p <* .001).

**Table 2.** Estimates of Linear Mixed Effects Model 1.1 for Planning Depth.

| | *b* | *SE b* | *p* | $\eta$ | *SE $\eta$* | *p* |
| --- | --- | --- | --- | --- | --- | --- |
| intercept | 1.758*** | .030 | < .001 | .027*** | .005 | < .001 |
| group<sup>a</sup> | .069 | .043 | .114 | - | - | - |
| steps | .710*** | .039 | < .001 | .044*** | .008 | < .001 |
| group x steps | .106 | .055 | .061 | - | - | - |

**Note.** *b* = fixed effect parameter; $\eta$ = random effect parameter; *SE* = standard error.
<sup>a</sup> the group variable was coded 0 / 1 for HC / AUD participants.

### Cognitive covariates

In terms of cognitive covariates, we did not observe performance differences between groups for the IDP task (*t*(60) = 1.02, *p =* .310) or the RAV task (*Z* = 1.89, *p =* .058). However, we observed significantly lower performance in HC participants in the SWM task (*t*(60) = 2.41, *p =* .019).

To control for the potentially confounding effect of cognitive covariates on task outcomes planning depth and performance, we included RAV- and SWM task performance in the linear mixed-effects model of planning depth (Model 1.2; see Table 3) as they correlated significantly with planning depth. This improved the model fit relative to model 1.1 (*χ^2^*(3) = 11.73, *p =* .008). In model 1.2, we observed significant main effects for steps (*t*(62.4) = 18.50, *p =* .007) and SWM performance (*t*(61.9) = 2.23, *p =* .029). There was no significant effect for group (*t*(62.4) = .59, *p =* .560), the group x steps interaction (*t*(62.3) = 1.92, *p =* .060), RAV performance (*t*(61.9) = 1.72, *p =* .091) or SAT RT (*t*(6175.9) = .24, *p =* .812).

**Table 3.** Estimates of Linear Mixed Effects Model 1.2 for Planning Depth.

| | *b* | *SE b* | *p* | $\eta$ | *SE $\eta$* | *p* |
| --- | --- | --- | --- | --- | --- | --- |
| intercept | .940** | .301 | .003 | .023*** | .004 | < .001 |
| group<sup>a</sup> | .024 | .041 | .560 | - | - | - |
| steps | .709*** | .038 | < .001 | .043*** | .008 | < .001 |
| group x steps | .106 | .055 | .060 | - | - | - |
| SAT_RT | .000 | .000 | .812 | - | - | - |
| SWM_PER | .008* | .004 | .029 | - | - | - |
| RAV_PER | .002 | .001 | .091 | - | - | - |

**Note.** *b* = fixed effect parameter; $\eta$ = random effect parameter; *SE* = standard error; SAT\_RT = Space Adventure Task reaction time [s]; SWM\_PER = Spatial Working Memory Task performance [%]; RAV\_PER = Raven Task Performance [%].
<sup>a</sup> the group variable was coded 0 / 1 for HC / AUD participants.

Secondly, we also included covariates IDP-, RAV- and SWM performance as well as SAT reaction time and a group indicator in a linear regression analysis of SAT performance. This revealed a significant effect for SAT reaction time (*b* = 1.163; *p* = .001), while the difference between groups was no longer significant (*b* = 4.724; *p* = .118; see Table S1).

### Model Parameters

Turning to the inferred parameters for the planning model, we observed significantly lower *β* values for HC participants (*t*(46.3) = 2.82, *p =* .007), indicating higher decision noise or potentially lower choice consistency. Further, Participants with AUD showed lower values of the response bias parameter 𝜃 (*t*(60) = -2.17, *p =* .034) meaning that they showed a stronger bias toward the deterministic “move” action. For the learning rate parameter 𝛼, HC participants showed higher values than Participants with AUD (*t*(31) = -6.15, *p <* .001), indicating a larger influence of recently observed transitions on the estimated jump success probability for HC. For parameter distributions across groups, see Figure 4.

**Figure 4.**
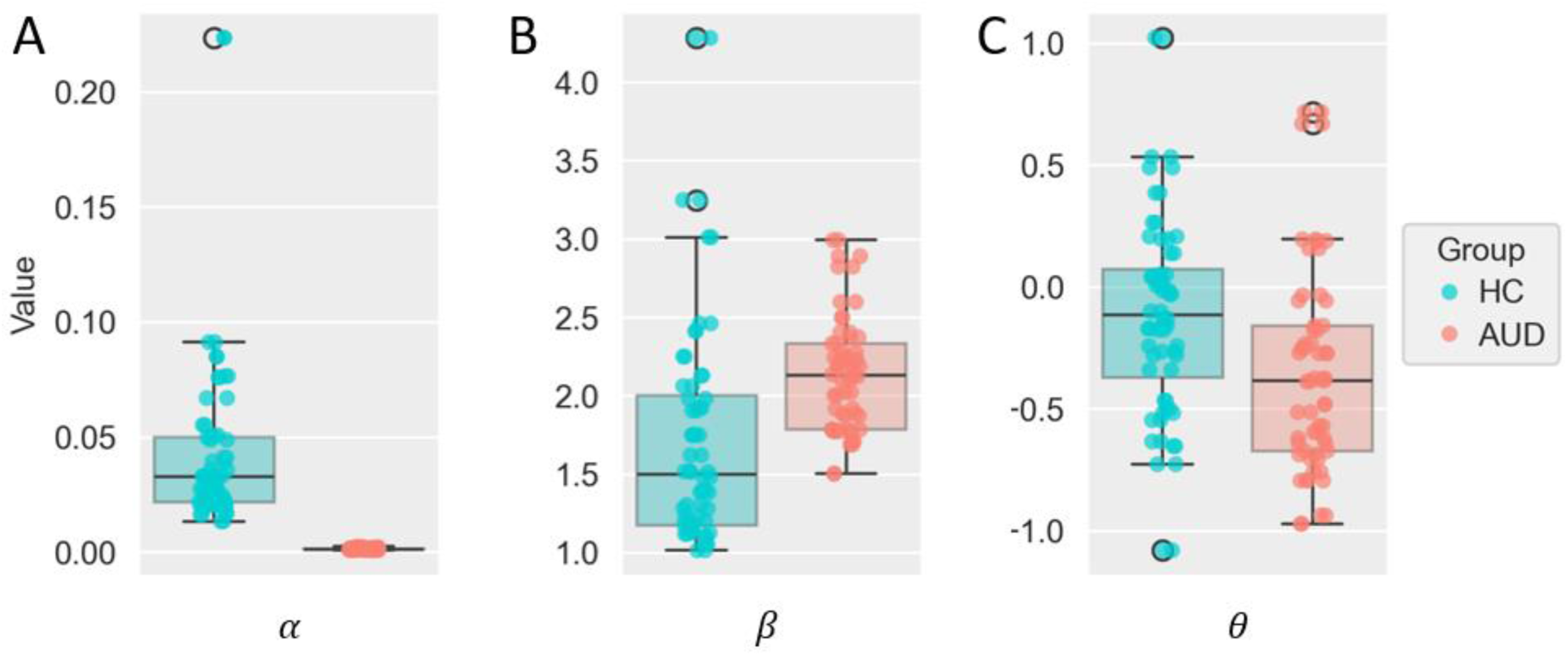
Inferred model parameter values per participant for the planning agent model in the Space Adventure Task. **(A)** Boxplot and scatterplot of participants’ inferred values for learning rate 𝛼 comparing healthy controls (HC) and participants with alcohol use disorder (AUD). **(B)** Same plot as (A) with values for inverse response noise *β*. **(C)** Same plot as (A) with values for response bias 𝜃.

## DISCUSSION

The current study was aimed at assessing forward planning in a general-population non-treatment-seeking sample with AUD in comparison to a HC group. As previous studies only focused on detoxified patients with long histories of alcohol-related problems or healthy samples without current problems, the intermediate stage during the development of AUD which our study focused on is not well characterized yet. A guiding hypothesis for our work has been that during this stage critical aspects of decision-making and forward planning lead to initial alcohol-related issues. Contrary to our expectations however, we did not find any evidence of lower forward planning capacity in individuals with mostly mild to moderate AUD. Indeed, we even found small effects in the opposite direction: the AUD group showed a significantly higher performance, spent more time planning and showed a slightly higher planning depth in the SAT, where the latter was marginally significant. Controlling for reaction time and general cognitive performances mostly eliminated between-group variance in planning depth and SAT performance. Moreover, while the AUD group showed longer reaction times in the SAT, they also showed a lower association of SAT reaction time with SAT performance.

Considering time as a valuable resource in the context of resource-rational accounts of cognitive control (Gershman et al., 2015; Shenhav et al., 2013), differences in SAT reaction time indicate that not only ability but also motivation may have driven the between-group differences in the SAT, where the AUD group possibly put, on average, more effort in the task. The post-hoc analysis of the effect of absolute Q-values of task trials on planning depths suggests that the incentive value of forward planning may have been a stronger motivational driver in the AUD group compared to healthy controls. Moreover, although mean performances in the spatial working memory task were in both groups comparable to previously reported accuracies in healthy participants (Nagel et al., 2008), we found higher performances in participants with AUD which could be another indicator of higher motivation.

A challenging theoretical property of forward planning tasks is that with deeper planning the computational costs typically increase exponentially. In an environment with relatively stable ranges of rewards, the ratio between computational costs and reward magnitude naturally decreases with increasing planning depth. Hence, forward planning tasks are particularly sensitive to motivational differences. Participants with AUD may have been more willing to accept a less efficient reward-computation-ratio and invest more planning time to increase their reward leading to higher performances in the SAT. However, as participants were explicitly instructed to “plan ahead to maximize reward”, we do not think that this tendency indicates a long-term liability. We rather speculate that boosted motivation for effort acts as a compensatory mechanism in individuals experiencing problems while not seeking treatment and still maintaining acceptable levels of functioning (Scheurich, 2005). This mechanism could also contribute to the high spontaneous remission rate of mild-to-moderate alcohol use disorder in the general population as those individuals might still get motivated to change their life-style after perceiving first serious consequences of their drinking behavior acting as punishments (Sobell et al., 2000; Tuithof et al., 2016).

A potential complication of our work could be a selection bias, i.e. resilient individuals who are coping well with minor problems in controlling their alcohol consumption might be more likely to participate in a study as ours (Tuithof et al., 2016). This group of people might also be particularly interested in getting accurate feedback on their current state of health and therefore be distinctively motivated to show good performance in the study whereas HC participants may be mainly motivated by monetary compensation.

### Limitations

A limitation of our work is that the task design turned out to be suboptimal to evaluate the effect of the noise condition, which therefore did not allow any interpretation. Moreover, the outbreak of the Covid-19 pandemic and the subsequent lockdown interrupted the assessment. In particular, while most control participants were assessed before the lockdown, most participants with AUD were assessed after the first lockdown, which took place from March to May 2020. This may have confounded our group comparison, which we could not account for. Furthermore, the cross-sectional nature of the current study was not suitable to be related to findings of previous longitudinal studies (Chen et al., 2021; Voon et al., 2015) pointing out the significance of future longitudinal investigations.

Finally, a potential true effect in the population may be too small to be detectable with a sample size as ours. Post-hoc sensitivity analysis with the HC group and only the participants with mild-to-moderate AUD showed that the minimum detectable population effect size with a power of 0.8 is 0.74.

## Conclusion

Our findings suggest that planning capacity, a key aspect underlying goal-directed behavior, is not reduced at this early stage of AUD. Conversely, our results suggest higher motivation to exert planning effort in the AUD sample. Further studies with larger sample sizes and longitudinal assessments of individuals with various AUD severities are needed to disentangle the dynamical interplay of forward planning, motivation and trajectories of AUD development.

## AUTHOR CONTRIBUTION STATEMENT

M.N.S. and S.J.K. conceptualized the study and supervised the study execution and data analysis. J.S. and P.C.F. contributed to the study design, implemented the study and conducted the assessments. J.S., L.G. and P.C.F. contributed to data analysis and to drafting the preprint version of the manuscript. J.S., L.G. and M.N.S. interpreted the results of the data analysis. J.S. implemented and performed the parameter inference and the final data analysis. J.S. wrote the manuscript and designed the plots. All authors reviewed the manuscript.

## DATA AVAILABILITY

All data generated or analyzed during this study along with the code and scripts necessary to perform the model-based inference and statistical analyses are available in the plandepth_aud Github repository, https://github.com/jeffensen/plandepth_aud.

