## Supplementary Material for "Forward planning in a population-based alcohol use disorder sample"

### SUPPLEMENTARY INFORMATION

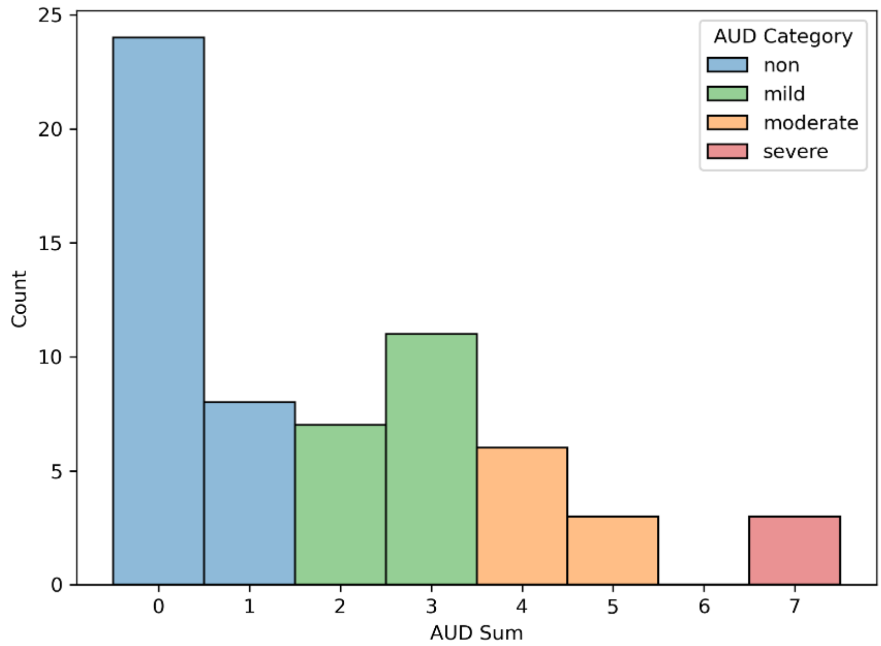


***Figure S1. Distribution of the sum of fulfilled criteria for alcohol use disorder (AUD) per participant.***

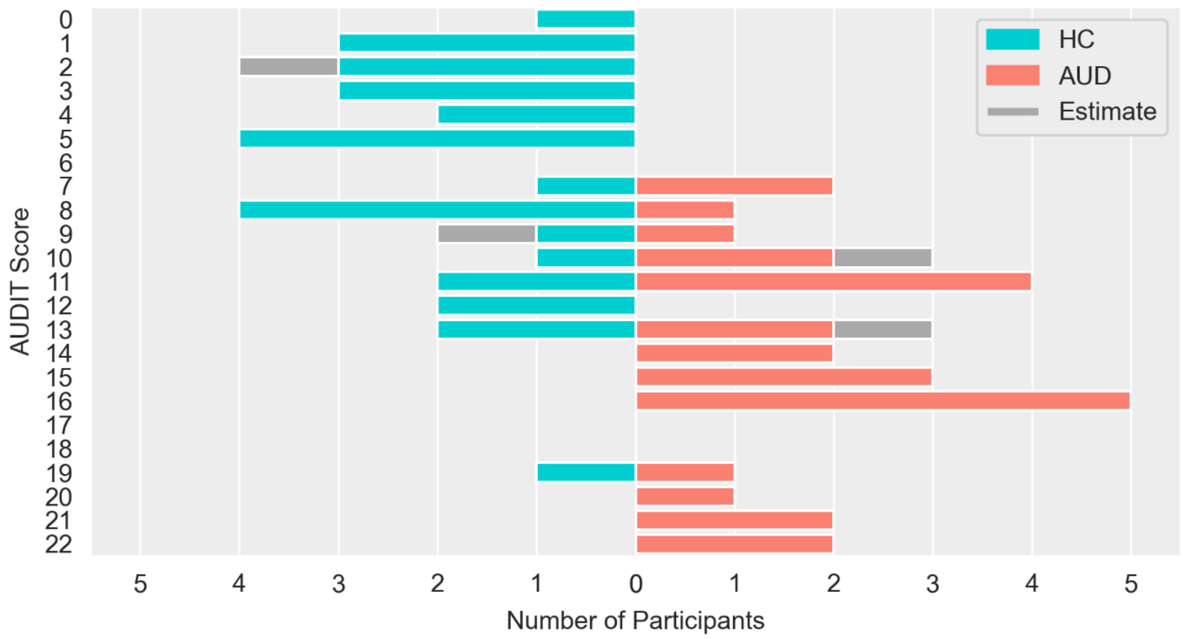


***Figure S2. Distribution of AUDIT scores per group.*** *Data was missing for four participants. Estimated values based on a telephone screening and SCID-AUD interview are added in grey ("Estimate").*
*AUDIT = Alcohol Use Disorders Identification Test.*

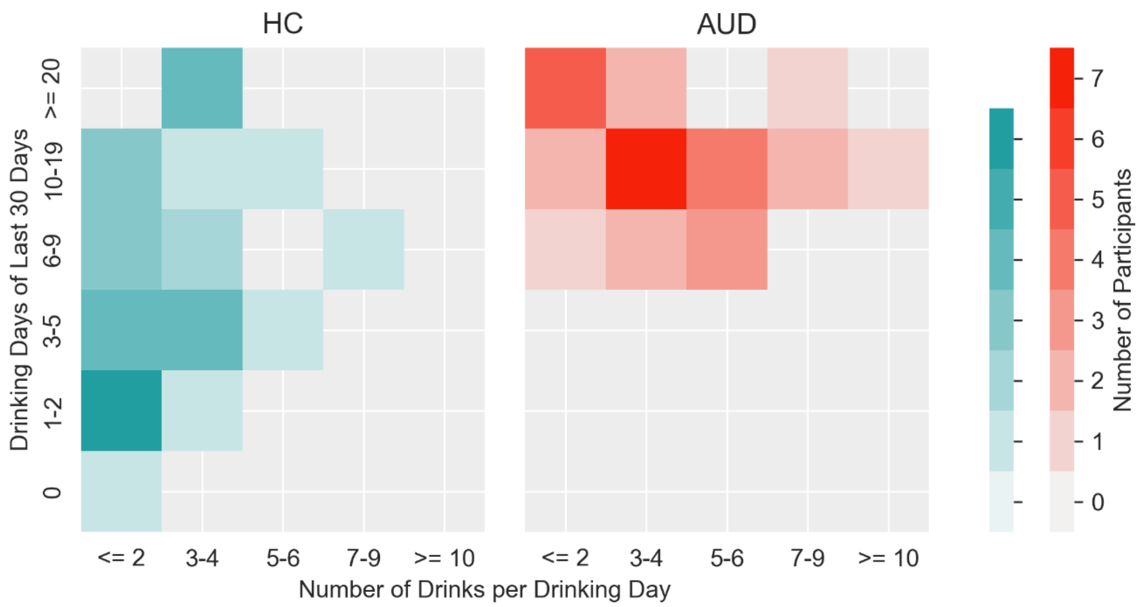


***Figure S3. Distribution of self-reported alcohol drinking frequency and quantity of last 30 days.*** *Values reflect participants categorical statements in the European School Survey Project on Alcohol and Other Drugs questionnaire (ESPAD) for last 30 days. However, we later replaced the ESPAD with a more valid quantity-frequency questionnaire (QF) leading to later participants making numerical statements for the last 3 months instead ($N_{HC}$=7, $N_{AUD}$=17). These numbers were therefore divided by three and assigned to the corresponding categories of the ESPAD. We thereby assumed that substance use during past 3 months was relatively constant.*

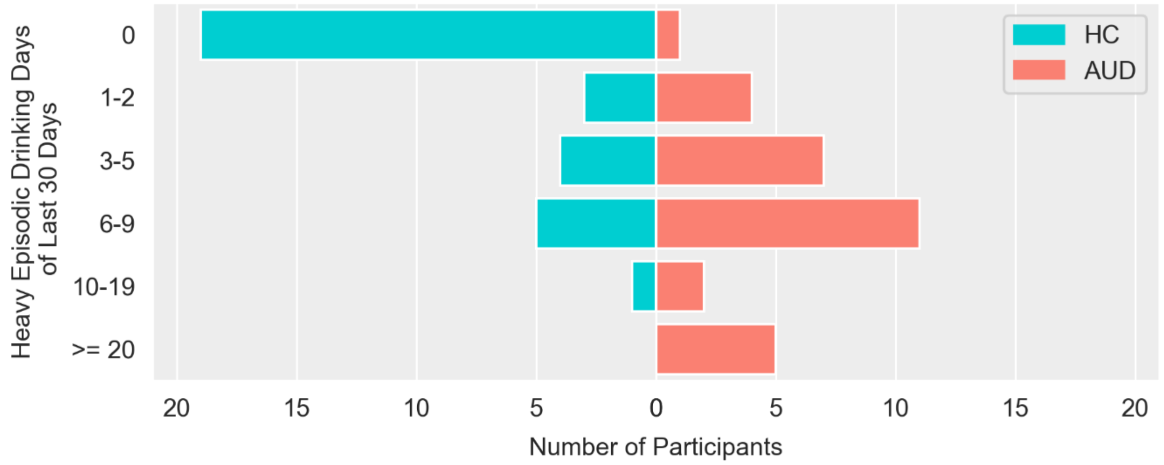


***Figure S4. Distribution of self-reported heavy episodic alcohol drinking frequency of last 30 days.*** *Heavy episodic drinking was defined as drinking >= 5 drinks on one occasion.*

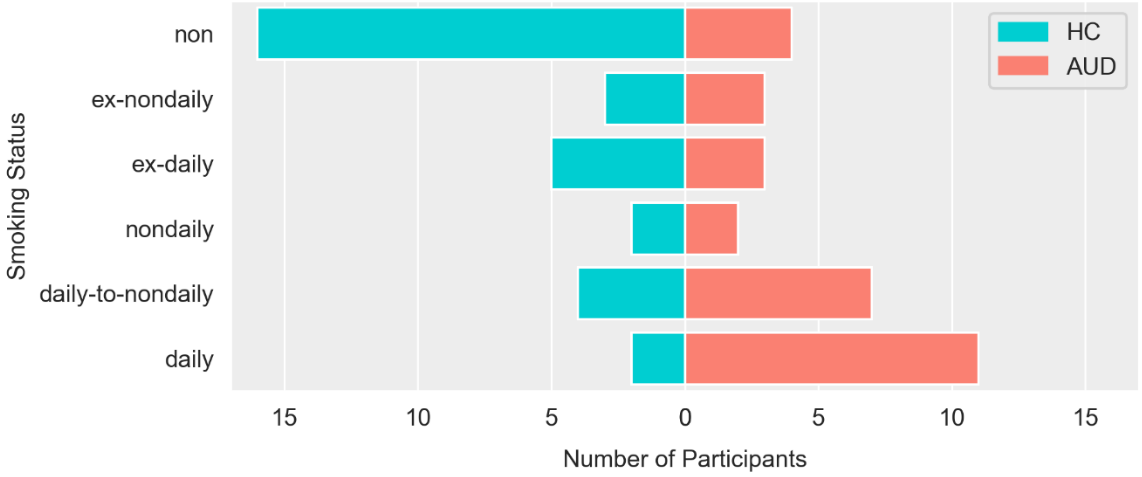


***Figure S5. Distribution of self-reported tobacco smoking status.*** *Categories refer to tobacco use of last 30 days with additional categories if lifetime maximum frequency was higher. The categories ex-(non)daily denote a current reduction from lifetime (non)daily smoking to no smoking during last 30 days. The category daily-to-nondaily denotes a current reduction from lifetime daily smoking to nondaily smoking during last 30 days.*

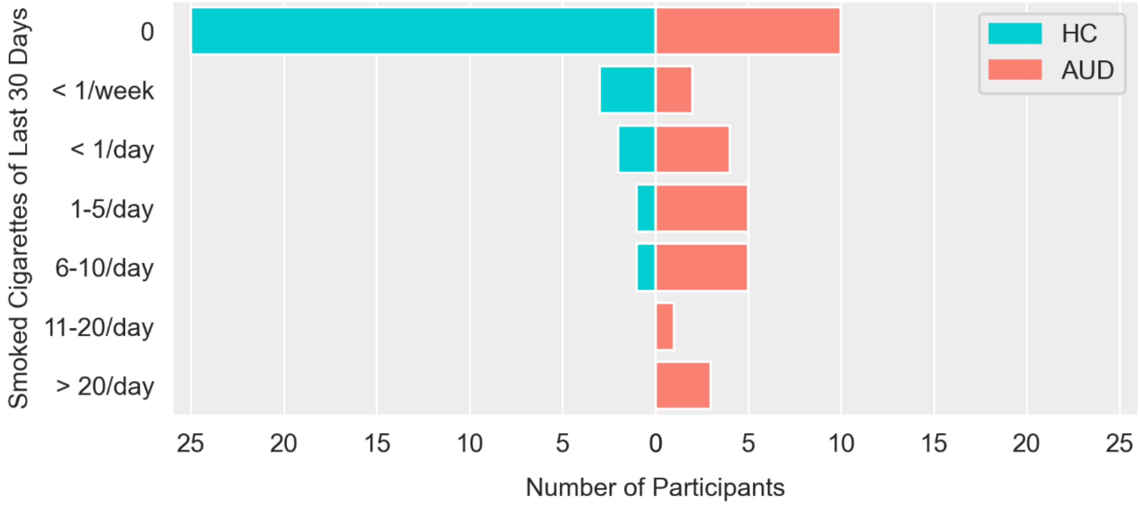


***Figure S6. Distribution of self-reported tobacco smoking quantity of last 30 days.***

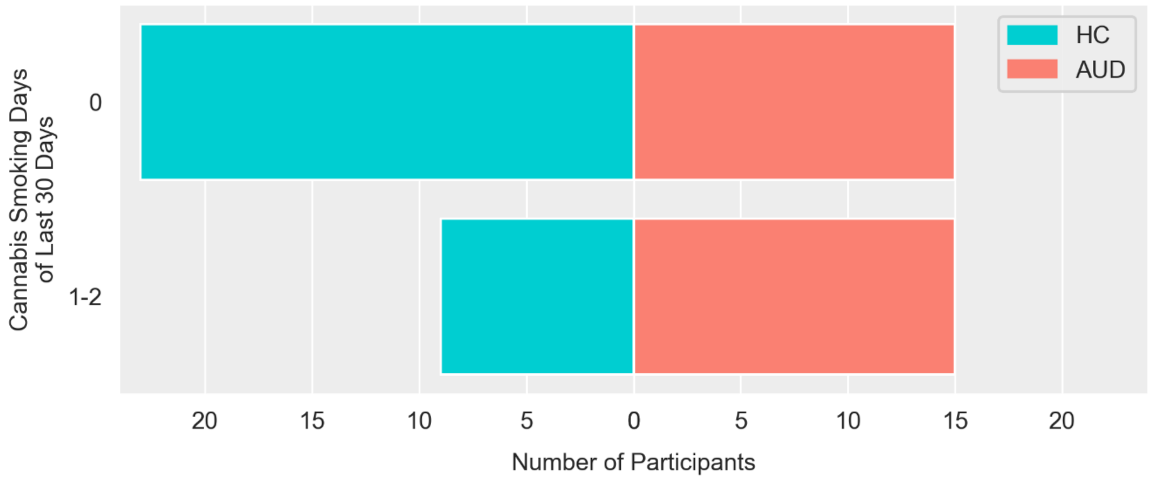


***Figure S7. Distribution of cannabis smoking frequency of last 30 days.***

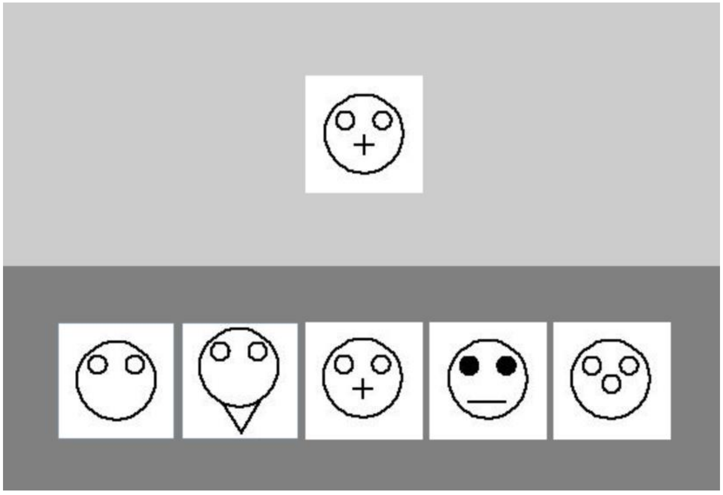

***Figure S8. Example trial of the Identical Pictures Task.*** *Participants were instructed to find out of 5 symbols the stimulus matching the target stimulus presented at the top by pressing the corresponding number on the keyboard (1-5) as quickly and accurately as possible. The task had an overall time limit of 80 seconds and performance was calculated as percentage of correct responses relative to the total of 46 trials.*

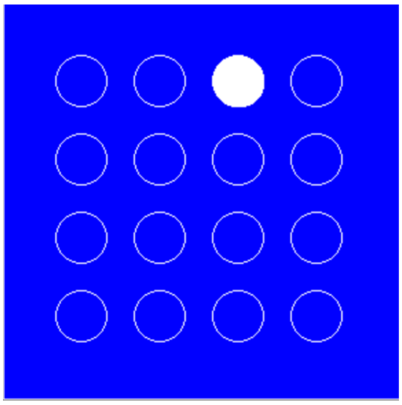

***Figure S9. Example stimulus of the Spatial Working Memory Task.*** *Participants were presented with a 4 x 4 grid of circles in which a series of dots were displayed consecutively in specific locations of the grid. At the end of each sequence, one circle was marked and participants had to indicate whether a dot was presented at that position or not (i.e., location memory condition). If they affirmed, a digit was shown at the marked position and participants then had to decide whether the dot was presented in that serial position or not (i.e., sequence memory condition). Working memory load was varied by dot sequence length either being 4 or 7. (i.e., working memory load levels). The time limit for each trial was 5 seconds for both conditions. As data for the sequence memory condition was very limited, we only used the location memory data for analysis and calculated performance as percentage of correct responses relative to the total of 96 trials across both load level.*

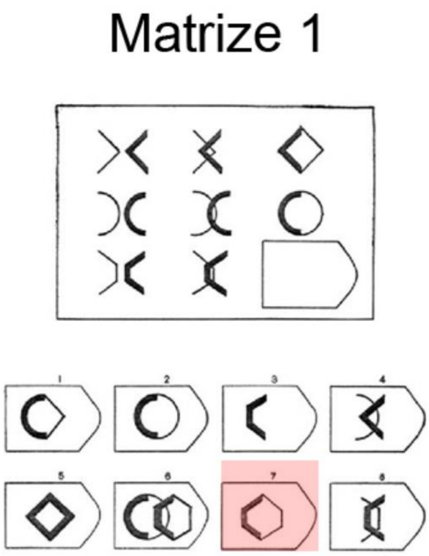


***Figure S10. Example trial of Raven's Matrices.*** *Participants were presented 12 matrices, each containing a 3 x 3 grid of shapes changing vertically and horizontally according to hidden logical rules. For each matrix ("Matrize" in German), the bottom-right shape was missing and participants had to choose the correct shape out of 8 options. There was a time limit of 15 minutes during which participants could freely navigate between matrices and change responses.*

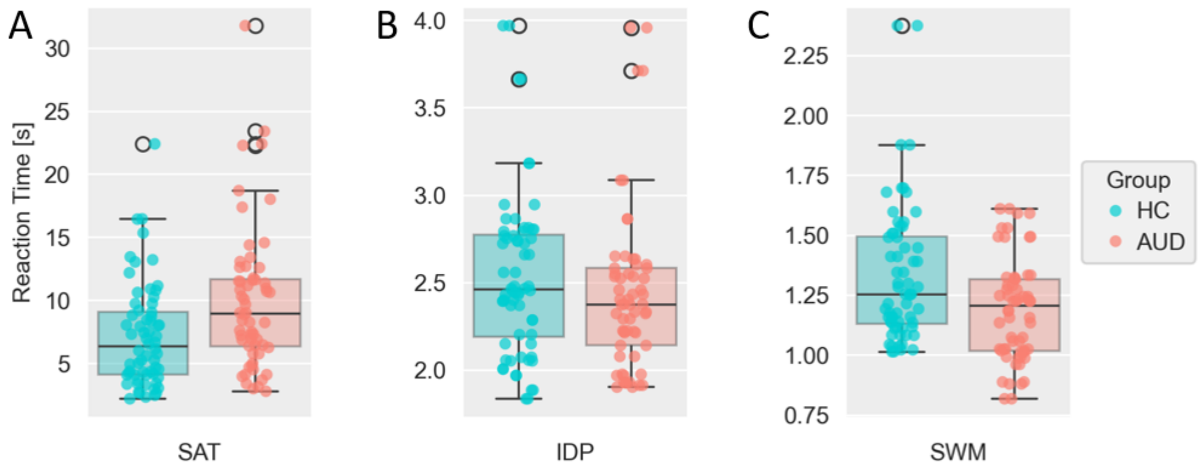


***Figure S11. Mean reaction times per participant of Space Adventure Task (SAT) and cognitive covariate tasks.*** ***(A)*** *Boxplot and scatterplot of participants' reaction times until execution of first action in the SAT comparing healthy controls (HC) and participants with alcohol use disorder (AUD).* ***(B)*** *Same plot as (A) with reaction times in the Identical Pictures Task (IDP).* ***(C)*** *Same plot as (A) with reaction times in the Spatial Working Memory Task (SWM).*

***Table S1 Estimates of Linear Regression Model for performance in the Space Adventure Task (SAT).***

| | *b* | *SE b* | *Beta* | *t* | *p* |
|---|---|---|---|---|---|
| intercept | 6.930 | 21.480 | | .323 | .748 |
| group[a] | 4.724 | 2.977 | .171 | 1.587 | .118 |
| SAT_RT | 1.163** | .342 | .389 | 3.399 | .001 |
| IDP_PER | -.134 | .162 | -.096 | -.833 | .409 |
| SWM_PER | .493 | .261 | .225 | 1.887 | .064 |
| RAV_PER | .114 | .079 | .169 | 1.434 | .157 |

***Note.*** *b = unstandardized coefficient; Beta = standardized coefficient; SAT_RT = Space Adventure Task reaction time [s]; IDP_PER = Identical Picture Task Performance [%]; SWM_PER = Spatial Working Memory Task performance [%]; RAV_PER = Raven Task Performance [%].*

*[a] the group variable was coded 0 / 1 for HC / AUD participants.*

***Table S2 Estimates of Linear Regression Model for mean planning depth in the Space Adventure Task (SAT).***

| | *b* | *SE b* | *Beta* | *Bootstrap*[b] | | |
|---|---|---|---|---|---|---|
| | | | | *p* | *95%CI* | |
| | | | | | *LL* | *UL* |
| intercept | 1.867*** | .071 | - | < .001 | 1.661 | 2.039 |
| group[a] | .239 | .106 | .540 | .071 | .003 | .503 |
| SAT_RT | .034** | .009 | .718 | .002 | .019 | .060 |
| SAT_RT*group[a] | -.021 | .011 | -.582 | .084 | -.054 | .003 |

***Note.*** *b = unstandardized coefficient; Beta = standardized coefficient; SAT_RT = Space Adventure Task reaction time [s].*
*[a] the group variable was coded 0 / 1 for HC / AUD participants.*
*[b] as the independent variable was non-normally distributed, we relied on non-parametric estimates using the bias-corrected and accelerated bootstrap approach with 10.000 samples (Efron & Tibshirani, 1994).*

***Table S3 Estimates of Linear Regression Model for performance in the Space Adventure Task (SAT).***

| | *b* | *SE b* | *Beta* | *t* | *p* |
|---|---|---|---|---|---|
| intercept | 38.559*** | 3.887 | - | 9.920 | < .001 |
| group[a] | 23.124*** | 5.753 | .836 | 4.019 | < .001 |
| SAT_RT | 2.743*** | .478 | .918 | 5.737 | < .001 |
| SAT_RT*group[a] | -2.047** | .613 | -.894 | -3.342 | .001 |

***Note.*** *b = unstandardized coefficient; Beta = standardized coefficient; SAT_RT = Space Adventure Task reaction time [s].*
*[a] the group variable was coded 0 / 1 for HC / AUD participants.*

***Table S4 Estimates of Linear Mixed Effects Model 1.3 for Planning Depth.***

| | *b* | *SE b* | *p* | *η* | *SE η* | *p* |
|---|---|---|---|---|---|---|
| intercept | 1.715*** | .037 | < .001 | .031*** | .007 | < .001 |
| group[a] | .028 | .054 | .606 | - | - | - |
| Q* | .018*** | .001 | < .001 | .000* | .000 | .014 |
| group x Q* | .004** | .002 | .008 | - | - | - |

***Note.*** *b = fixed effect parameter; η = random effect parameter with variation by participant; SE = standard error; Q* = maximum of absolute Q values per mini-block of all possible planning depths and actions.*
*[a] the group variable was coded 0 / 1 for HC / AUD participants.*

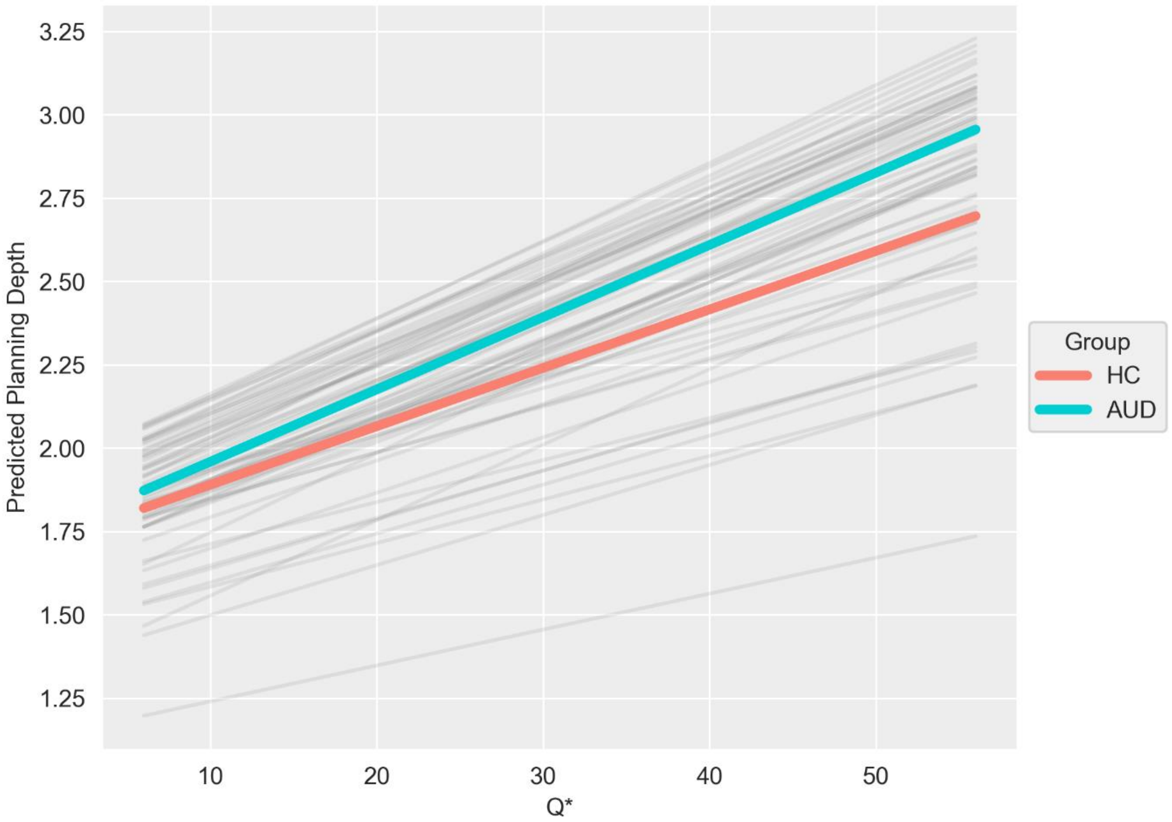


***Figure S12. Lineplot of mean planning depths predicted by linear mixed-effects model 1.3 and maximum absolute Q-values (Q*).*** *The latter served as an indicator for incentive value of each mini-block. For this, we first computed Q-values for each mini-block, action and planning depth and extracted the maximum absolute value for each mini-block. Grey lines indicate the association per participant.*

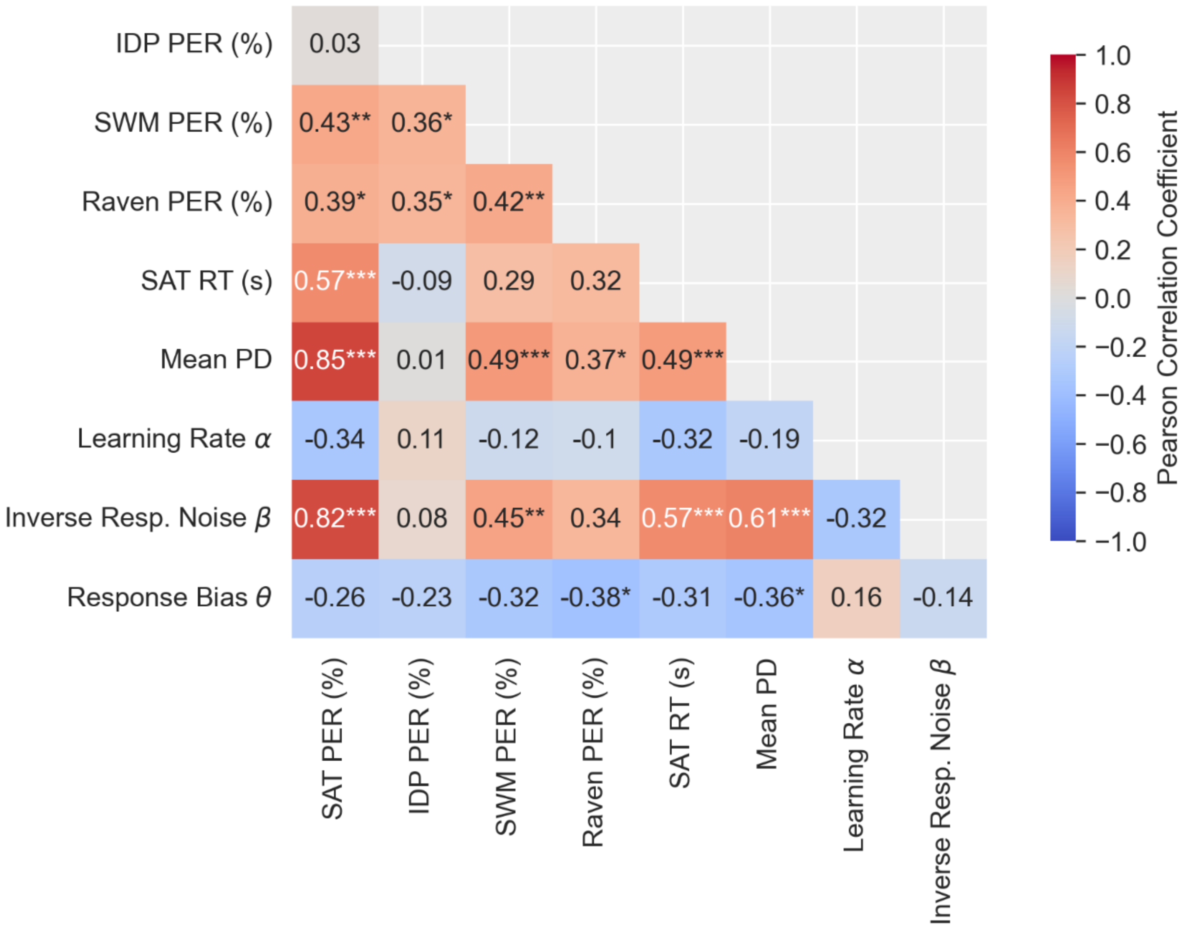


***Figure S13. Heatmap of Pearson correlations of all task outcomes.***

** / ** / *** significant at alpha = .05 / .01 / .001 (Bonferroni-Holm corrected).*

*SAT = Space Adventure Task. IDP = Identical Pictures Task. SWM = Spatial Working Memory Task. PER = Performance. RT = Reaction Time. PD = Planning Depth.*
